# Differential Effects of Neutral Sphingomyelinase and Serine Palmitoyltransferase Inhibitors on Alzheimer’s-Like Neuropathology

**DOI:** 10.64898/2026.09.04.749232

**Authors:** Melina P. Bellotto, Nicolás González Pérez, Melisa Bentivegna, Martín Carrera, Jessica Presa, Angeles Vinuesa, Carlos Pomilio, Flavia Saravia, Juan Beauquis

**Affiliations:** Neurobiology of Aging Laboratory, Institute of Biology and Experimental Medicine (IBYME), National Research Council, Buenos Aires, Argentina; Department of Biological Chemistry, School of Exact and Natural Sciences, University of Buenos Aires, Argentina

## Abstract

Alzheimer’s disease (AD) involves progressive neurodegeneration, amyloid-β (Aβ) pathology and chronic neuroinflammation. Elevated ceramides, generated *via* neutral sphingomyelinase (nSMase)-mediated sphingomyelin hydrolysis or serine palmitoyltransferase (SPT)-driven *de novo* synthesis, amplify these processes. Here, we performed a direct comparison of pharmacological inhibition of these pathways in the PDAPP-J20 transgenic mouse model and in fibrillized Aβ1-42-challenged BV-2 murine microglial cell line.

Eight-month-old PDAPP-J20 female mice received intraperitoneal GW4869 (nSMase inhibitor, 1.25 mg/kg) or myriocin (SPT inhibitor, 0.3 mg/kg) three times weekly for 3 weeks. Neutral SMase inhibition restored spatial learning in the Barnes maze, reduced hippocampal neuronal loss and layer atrophy, decreased amyloid plaque burden, and attenuated microglial activation (Iba1 morphology and peri-plaque reactivity). In contrast, SPT inhibition worsened thigmotaxis, failed to improve cognition, and increased plaque load.

*In vitro*, nSMase blockade (GW4869 and cambinol) suppressed Aβ-induced NFκB p65 nuclear translocation, blunted TNF-α expression, and reduced intracellular Aβ accumulation in microglia, suggesting enhanced endolysosomal degradation. SPT inhibition lacked these anti-inflammatory and clearance-promoting effects.

These results identify nSMase as a critical node linking ceramide metabolism, microglial dysfunction, and amyloid progression in AD. Targeted nSMase inhibition offers a promising molecular strategy to interrupt neuroinflammation and restore neuronal homeostasis, while broad SPT blockade appears counterproductive. Pathway-selective modulation of ceramide signaling may open new therapeutic avenues for AD.

**Highlights:**

- nSMase inhibition restores learning and reduces hippocampal damage in AD mice.
- GW4869 decreases amyloid plaque burden and attenuates microglial activation.
- SPT inhibition worsens anxiety-like behavior and increases hippocampal plaque load.
- nSMase blockade reduces NFκB/TNFα signaling and intracellular Aβ in BV-2 microglia.
- Pathway-specific ceramide modulation is a promising therapeutic target in AD.

## Introduction

Alzheimer’s disease (AD) is characterized by progressive neurodegeneration, amyloid-β (Aβ) plaques, tau pathology, and neuroinflammation. Although the classical hallmarks of AD center on amyloid and tau pathologies, lipid dyshomeostasis has emerged in recent years as a key contributor to disease progression, offering promising new therapeutic targets. Elevated ceramide levels have been consistently observed in AD brains (particularly long-chain species such as Cer16, Cer18, Cer20, and Cer24), as well as in cerebrospinal fluid (CSF) and plasma of affected individuals, correlating with cognitive decline, hippocampal atrophy, and neuropathological burden (Czubowicz et al., 2019; Filippov et al., 2012; Mielke et al., 2010; Teitsdottir et al., 2021; van Kruining et al., 2023; Vashishath et al., 2026). These bioactive sphingolipids regulate fundamental processes including apoptosis, oxidative stress, inflammation, and endosomal trafficking, contributing to neuronal damage and Aβ accumulation (Kalkman and Smigielski, 2025; Pan et al., 2024).

Ceramides are generated through multiple pathways, including *de novo* synthesis *via* serine palmitoyltransferase (SPT) and hydrolysis of sphingomyelin by neutral sphingomyelinase (nSMase). Neutral SMase, activated by pro-inflammatory cytokines such as TNFα, triggers NFκB signaling and promotes cytokine expression, amplifying glial reactivity (Al-Rashed et al., 2020; Huang et al., 2025; Trajkovic et al., 2008). Additionally, TNFα-mediated nSMase activation elevates plasma membrane ceramides, increasing membrane rigidity and stabilizing amyloid precursor protein (APP) processing machinery, thereby enhancing Aβ production and perpetuating a vicious cycle of inflammation and neurodegeneration (Kalkman and Smigielski, 2025; Tohumeken et al., 2023).

Given ceramides’ central role in linking inflammation, glial dysfunction, and amyloid pathology, we hypothesized that selective inhibition of ceramide synthesis pathways would modulate microglial responses in AD-like pathology, attenuating the amplification and propagation of inflammatory signals. To test this, we examined the effects of pharmacological inhibitors GW4869 (nSMase inhibitor) and myriocin (SPT inhibitor) in the PDAPP-J20 transgenic mouse model of AD and in fibrillized Aβ₁₋₄₂-challenged BV-2 microglia *in vitro*. These interventions were evaluated for impacts on neurodegeneration, amyloid burden, microglial activation, behavioral deficits, and molecular inflammatory markers.

## Materials and Methods

### Animals and Treatments

Adult female PDAPP-J20 transgenic mice (B6.Cg-Zbtb20Tg(PDGFB-APPSwInd)20Lms), hemizygously expressing human amyloid precursor protein (APP) with the Swedish (K670N/M671L) and Indiana (V717F) mutations under the PDGF-β promoter (Mucke et al., 2000), were obtained from the Institute of Biology and Experimental Medicine Animal Facility (NIH Assurance Certificate #A5072-01) and genotyped by PCR as previously published (Bentivegna et al., 2025). These mice exhibit Aβ plaque deposition and cognitive deficits starting from approximately 5–6 months of age (Beauquis et al., 2014; Pomilio et al., 2016). Non-transgenic littermates served as controls (CTL).

Eight-month-old female mice were randomly assigned to four experimental groups (n ≥ 10 per group): CTL (non-transgenic-vehicle), TG (transgenic-vehicle), TG-GW4869, and TG-myriocin (Myr). Non-transgenic mice also were treated with the drugs but their performance in behavioral tests was not different from vehicle-treated mice and thus not included in the comparisons (see suppl. material). Treatments were administered *via* intraperitoneal (i.p.) injection (32 G needle, 100 µL per injection) three times per week for 3 weeks (total 9 injections). Mice were handled by the experimenter on the previous days to minimize stress during injections. GW4869 (Cayman Chemical, Ann Arbor, MI, USA) was dosed at 1.25 mg/kg, and Myr (Cayman Chemical) at 0.3 mg/kg. Doses were set following a preliminary pilot study and previous reports (Hojjati et al., 2005; Mowry et al., 2023; Tabatadze et al., 2010). No overt adverse effects (e.g., weight loss, abnormal behavior, poor coat state, etc.) were observed during treatment (suppl. fig. 1). The vehicle consisted of 25% DMSO in sterile saline.

At the end of the treatment period, mice were euthanized by anesthesia with Ketamine (0.85 mg/Kg) and Xylazine (0.1 mg/Kg) i.p. followed by decapitation, and brains were collected for histological and biochemical analyses. Right brain hemispheres were fixed, cryoprotected, and sectioned into 60-μm-thick serial coronal slices using a vibratome (PELCO easiSlicer, Ted Pella, USA) and stored in cryoprotectant solution (PBS-Glycerol-Ethylene glycol, 50:25:25 v/v/v) at – 20°C until use. Left brain hemispheres were dissected to obtain hippocampi that were snap frozen and kept at -80°C for future studies.

All animal procedures complied with the ARRIVE (Animal Research: Reporting of In Vivo Experiments) guidelines, were conducted in accordance with the NIH Guide for the Care and Use of Laboratory Animals and were approved by the Institutional Animal Care and Use Committee of the Institute of Biology and Experimental Medicine (protocol #20/2023). All efforts were done to minimize animal suffering and discomfort. A total of 57 mice were used in this study, maximizing the number of techniques performed per animal in order to reduce the total number of mice while keeping biological meaningfulness. Only female mice were used (see Discussion for rationale). Exclusion of animals for any experiment was based only on statistical (outliers) or technical reasons (mice for analysis were randomly selected).

### Behavioral Testing

Spatial learning was evaluated using the Barnes maze, adapted from Bentivegna et al. 2025 (Bentivegna et al., 2025). Briefly, mice were acclimatized for 30 min to the behavior testing room and subjected to 4 training trials per day for 4 days. Each trial consisted in letting mice explore the platform (white, ⌀ 85 cm, elevated 70 cm from the ground with 20 evenly distributed ⌀ 6 cm-holes in the periphery of the platform) until they found the escape hole that led to a safe box or until 90 s elapsed without finding it. On the fifth day, mice were tested in the platform with the escape box removed, allowing them to explore it for 90 s. Locomotor and anxiety-like behavior were assessed using the open field (OF) test, adapted from Vinuesa et al. 2016 (Vinuesa et al., 2016). Briefly, mice were allowed to explore the arena (white, 55 cm L × 55 cm W with 15 cm H walls) during 300 s, All sessions were video-recorded, coded and blindly analyzed using Any-Maze software (Stoelting Co., Wood Dale, IL, USA) to quantify parameters including total distance traveled, mean velocity, thigmotaxis (preference for the periphery and avoidance of the open center of the OF, a widely used measure of anxiety-like behavior; peripheral distance was defined as a 11 cm wide corridor along the walls of the OF arena), and escape latency and distance (time and distance traveled to find the target hole in the Barnes maze).

### Nissl Staining and Neuronal Quantification

Cryoprotectant was removed from selected sections by PBS washes. Sections were mounted on gelatin-coated slides, air-dried overnight, and stained with 0.1% cresyl violet for 20 min at room temperature. Dehydration and clearing were performed in graded ethanol (95% to 100%) and xylene series. Slides were coverslipped with Canada Balsam (Biopack, Argentina). Bright-field images were captured using a Nikon E200 microscope with a Micrometrics 318CU CMOS camera and 40× air objective. Neuronal density and pyramidal/granular layer thickness in CA1 and dentate gyrus were quantified blinded in FIJI/ImageJ (Schindelin et al., 2012).

### Congo Red Staining and Amyloid Quantification

Amyloid pathology was assessed only in transgenic groups (n = 5 per treatment). Brain sections containing hippocampi were washed in PBS, stained with 0.1% Congo red in 85% ethanol/NaOH (4%) for 5 min, washed in PBS, dehydrated in graded ethanol and xylene, and mounted with Canada Balsam. Imaging and quantification of plaque area and number in hippocampal regions *stratum radiatum* under CA1 and dentate gyrus were performed blinded using FIJI/ImageJ on bright-field images (Nikon E200, Micrometrics 318CU CMOS camera, 40× air objective).

### Immunohistochemistry and Confocal Imaging

For each mouse, six representative hippocampal sections were selected (n = 5 mice per group).

Free-floating immunostaining was performed as follows: sections were washed in PBS, blocked in PBS containing 0.1% Triton X-100 and 1% normal goat serum, and incubated overnight at 4°C with primary antibodies diluted in blocking solution. After PBS washes, sections were incubated with appropriate fluorescent secondary antibodies for 2 h at room temperature, counterstained with DAPI (1:5000; Sigma-Aldrich, St. Louis, MO, USA) for 10 min, and mounted with PVA-DABCO (Sigma-Aldrich). Primary antibodies included: anti-Iba-1 (1:500; 019-19741, Wako Chemicals, Japan), anti-4G8 (1:700; MAB1561-M, Millipore Sigma, USA). Confocal images were acquired using an Olympus IX83 DSU microscope and a 40× air objective. Quantification of the Iba1 activation score was done following previously published procedures (González Pérez et al., 2025), discriminating amyloid plaque-associated microglia (less than 25 µm from the plaque center) from cells not associated to plaques or in CTL mice. The radial profile analysis was done blindly using the Radial profile plugin for ImageJ by Paul Baggethun (Pittsburgh, PA).

### Cell Culture and In Vitro Treatments

The murine BV-2 microglial cell line was cultured in RPMI (Sigma-Aldrich) with 10% FBS (Internegocios, Argentina) and Penstrep (Sigma) at 37°C, 5% CO₂. Cells were seeded in 24 well-plates with ⌀ 12 mm coverslips and allowed to adhere overnight. Aβ₁₋₄₂ (Tocris) was fibrillized by incubation at 37°C for 3 days in sterile distilled water. Fibrillized Aβ1-42 (fAβ1-42) was added at 0.5 or 0.05 μM, as specified in each result, during 30 min for NFκB detection and 24 h for RT-qPCR. Treatments included: GW4869 (10 μM), Myr (1.5 µM), Cambinol (alternative nSMase inhibitor; 10 µM, Cayman Chemicals), or vehicle (DMSO 0.1%). Cells were pretreated with inhibitors for 2 h before fAβ1-42 exposure to allow effective inhibition of the enzymes (Camell et al., 2015; Kumar et al., 2019)

### Immunofluorescence for NFκB Nuclear Translocation

Following treatment, cells were fixed in 4% paraformaldehyde, permeabilized, blocked, and incubated with anti-NFκB p65 antibody (1:500; sc-372, Santa Cruz Biotechnology) overnight at 4°C, followed by fluorescent secondary antibody and DAPI counterstaining. Nuclear translocation was quantified by densitometric analysis using FIJI/ImageJ on blindly-coded confocal images obtained with an Olympus IX83 DSU microscope.

### Quantitative RT-PCR for TNF-α

Total RNA was extracted using TransZol (TransGen Biotech Co., China), reverse-transcribed (MMLV Reverse Transcriptase, Promega Corporation, USA), and qPCR performed with primers for TNF-α and GAPDH (IDT Inc., USA; Suppl. Table 1) using the FastStart Universal SYBR Green Master (Rox) mix (Roche) in a Bio-Rad CFX system (Biorad) following previously published protocols (Vinuesa et al., 2019). Relative expression was calculated by the 2(-ΔΔCt) method (Pfaffl, 2001).

### Phagocytosis Assay with TAMRA-Labeled Aβ

Cells were incubated with fibrillized and sonicated Aβ-TAMRA (0.05 μM, Cayman Chemicals) for 6 h ± inhibitors following previously published protocols (Zyśk et al., 2023). Intracellular fluorescence intensity and cell area were quantified using FIJI/ImageJ on blindly-coded confocal images (Olympus IX83 DSU, 40× air objective).

### Statistical Analysis

Statistical analyses were performed using GraphPad Prism version 9.0 (GraphPad Software, San Diego, CA, USA). Data were assessed for normality using the Shapiro–Wilk test. Normally-distributed data were analyzed using unpaired two-tailed Student’s t-tests, one-way ANOVA, or two-way ANOVA, followed by appropriate *post hoc* tests as detailed in each result. Outliers were detected using Grubb’s test. Reported p values were adjusted for multiple comparisons. In cases where sample sizes were too small to reliably assess normality or when data were non-normally distributed, Kruskal–Wallis test was applied, as detailed in each case.

All data are presented as mean ± standard error of the mean (SEM). The type and number of replicates (biological and/or technical) for each experiment are specified in the corresponding figure legends.

Sample sizes were prospectively determined using StatMate 2.0 (GraphPad Software) based on effect sizes and variability observed in our prior published studies. Calculations targeted a significance level of α = 0.05 and a power of 80% (1 – β = 0.80) to detect a minimum between-group difference of 10%. This approach balanced the detection of biologically meaningful effects with the ethical imperative to minimize animal use.

## Results

### GW4869 Restores Spatial Learning in the Barnes Maze and Differentially Modulates Locomotor and Anxiety-Like Behaviors in the Open Field in PDAPP-J20 Transgenic Mice

Adult PDAPP-J20 and control mice were treated for 3 weeks with GW4869 (GW) or myriocin (Myr), inhibitors of nSMase and SPT ceramide synthesis pathways, respectively. In the last week of the treatment, locomotor activity, anxiety-like behavior, and cognition were evaluated (fig. 1a). In the Barnes maze, spatial learning was assessed during the training phase by calculating the ratio of escape latency on the final trial of day 4 (T4-D4) to that on the final trial of day 1 (T4-D1), providing a normalized index of learning improvement across sessions. One-way ANOVA revealed a difference between groups (F (3,44) = 3.847, P < 0.05; fig. 1b), with transgenic (TG) mice exhibiting higher escape latency ratios (poorer learning performance) compared to non-transgenic controls (CTL; Tukey’s P < 0.05). GW treatment fully restored this learning parameter in TG mice (P < 0.05), making performance levels comparable to CTL. In contrast, Myr treatment provided no significant improvement over vehicle-treated TG mice. Twenty-four h after the last training trial, a memory test was conducted by removing the escape box and allowing mice to freely explore the platform for 90 s. Search strategies were qualitatively poorer in TG mice compared with CTL (fig. 1c). Differences were found in the latency (one-way ANOVA, F (3,46) = 6.661, P < 0.01) and distance (Kruskal-Wallis P < 0.05) required to reach the target (fig. 1d-e). Transgenic mice treated with GW showed lower latencies to the target compared with untreated TG (Holm-Sidak’s P < 0.05). Myriocin treatment did not show an effect on these parameters, suggesting that cognitive recovery was selectively associated with inhibition of the nSMase-dependent ceramide synthesis pathway. Neither GW nor Myr exerted significant effects on non-transgenic mice in these parameters (suppl. fig 2).

**Figure 1.**
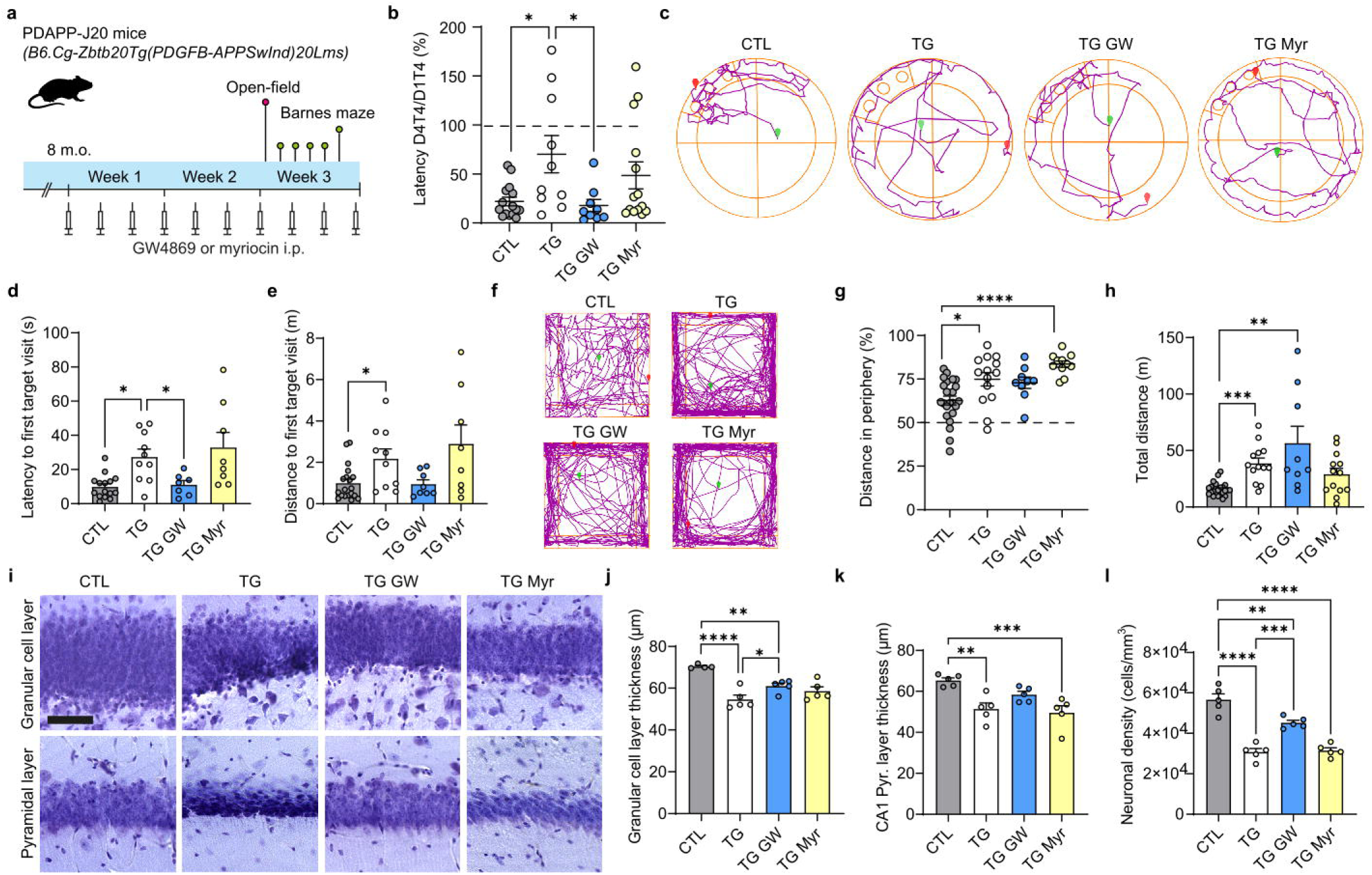
Experimental design, behavioral outcomes, and hippocampal histological analyses. **a.** *In vivo* experimental protocol. Adult 8-month-old PDAPP-J20 (TG) mice were treated i.p. for 3 weeks with GW4869 (GW) or myriocin (Myr) and then behaviorally evaluated in the open-field and Barnes maze tests. **b.** Escape latency ratio in the Barnes maze. For learning ability assessment, the ratio between escape latencies on the last trial of the 4^th^ training day and on the last trial of the 1^st^ training day (learning index D4T4/D1T4; CTL n=15, TG n=10, TG GW n=9, TG Myr n=14). **c.** Representative track plots in the test trial (5^th^ day) of the Barnes maze (generated by Anymaze software). Green and red symbols represent the start and end of mice trajectories during 90 s, respectively. **d and e.** Latency and distance to the target hole on the test trial (CTL n=15, TG n=10, TG GW n=8, TG Myr n=8). **f.** Representative track plots in the open-field for 5 min of exploration. **g and h.** Percentage of distance travelled in the periphery (thigmotaxis) and total distance travelled in the open-field for 5 min (CTL n=23, TG n=14, TG GW n=9, TG Myr n=11. **i.** Representative microphotographs of Nissl-stained hippocampal sections showing the granular cell layer (GCL) of the dentate gyrus (upper row) and the pyramidal layer of CA1 (lower row). Scale bar=40 µm. **j and k.** Thickness of the granular and CA1 pyramidal layers of the hippocampus. **l.** Neuronal density in the GCL measured on Nissl-stained sections (n=5 mice per group). All data is presented as the mean ± SEM. Statistical tests performed are described in the text. Asterisks represent significant differences found in the corresponding *post hoc* tests. * P < 0.05, ** P < 0.01, *** P < 0.005, **** P < 0.001.

In contrast to the selective cognitive benefit of nSMase inhibition, neither treatment normalized hyperlocomotion or thigmotaxis. In the open-field test (fig. 1f), mice showed differences on thigmotaxis (one-way ANOVA F (3,54) = 8.942, P < 0.005; fig. 1g) and total distance travelled (Kruskal-Wallis P < 0.001; fig. 1h). Relative to CTL, TG mice displayed increased thigmotaxis (Tukey’s P < 0.05) and total distance travelled (Dunn’s P < 0.005), suggesting anxiety-like behavior. Neither GW nor Myr significantly improved these parameters in TG mice. Moreover, TG mice treated with Myr showed a further increase in peripheral preference (P < 0.001 vs. CTL). Neither GW nor Myr exerted significant effects on the behavior of non-transgenic mice in the open-field test (suppl. fig 3).

### Neutral SMase Inhibition Reduces Hippocampal Neuronal Loss and Layer Atrophy in PDAPP-J20 Mice

After behavioral assessment, mice were euthanized and brains processed for histological analyses. To assess hippocampal neuronal integrity, brain sections were stained with the Nissl technique, enabling the evaluation of the granular cell layer (GCL) in the dentate gyrus and of the pyramidal layer in the CA1 region (fig. 1i). One-way ANOVA revealed differences in the thickness of both layers (GCL: F (3,15) = 14.28 P < 0.005; Pyr CA1: F (3,16) = 8,66 P < 0.001; fig. 1j-k). Vehicle-treated TG showed thinner layers compared to those of CTL mice (GCL: Sidak’s P < 0.001; Pyr CA1: Tukey’s P < 0.01) while GW4869 administration yielding an improvement in GCL thickness (TG GW vs. TG; Sidak’s P < 0.05). Neuronal density of the GCL was lower in TG vs. CTL mice (one-way ANOVA F (3,16) = 40.81 P < 0.001; Tukey’s P < 0.001; fig. 1l). This genotype-associated deficit was attenuated by treatment with GW4869 (TG GW vs. TG; P < 0.005) but not with Myr. Nevertheless, GCL neuronal density was lower in both TG GW and TG Myr when compared with CTL (P < 0.01 and 0.001, respectively).

### GW4869 Reduces Parenchymal Amyloid Burden in the Hippocampus of PDAPP - J20 Transgenic Mice

To evaluate hippocampal amyloid pathology, the core histopathological hallmark of AD, we performed a Congo red staining on brain sections of TG mice (fig. 2a) and measured hippocampal amyloid plaque load and number of plaques per section in the *stratum radiatum* under CA1 and in the dentate gyrus. The statistical analysis showed differences in the plaque load between groups in both regions (CA1: one-way ANOVA F (2,12) = 19.11 P < 0.005; dentate gyrus: Kruskal-Wallis P < 0.05; fig. 2b-c). Transgenic mice treated with GW4869 showed lower CA1 plaque load compared with both TG and TG Myr (Tukey’s P < 0.05 and 0.005, respectively; fig. 2b). Myriocin treatment was associated with higher CA1 plaque load compared with vehicle-treated TG (P < 0.05; fig. 2b), suggesting a worsening of hippocampal amyloid pathology with this inhibitor. Plaque load was also higher in the dentate gyrus of TG Myr compared to that of TG GW (Dunn’s P < 0.05; fig. 2c), though neither inhibitor induced differences with vehicle-treated TG. Regarding the number of plaques per hippocampal section, one-way ANOVA showed differences only in the dentate gyrus (CA1: F (2,13) = 3.311 P = 0.06; dentate gyrus: F (2,12) = 19,71 P < 0.005). Transgenic mice treated with GW showed a trend towards lower number of plaques (Sidak’s P = 0.064) while TG Myr showed an increase in this parameter compared with TG (P < 0.01), confirming a worsening of amyloid pathology seen in the plaque load assessment.

**Figure 2.**
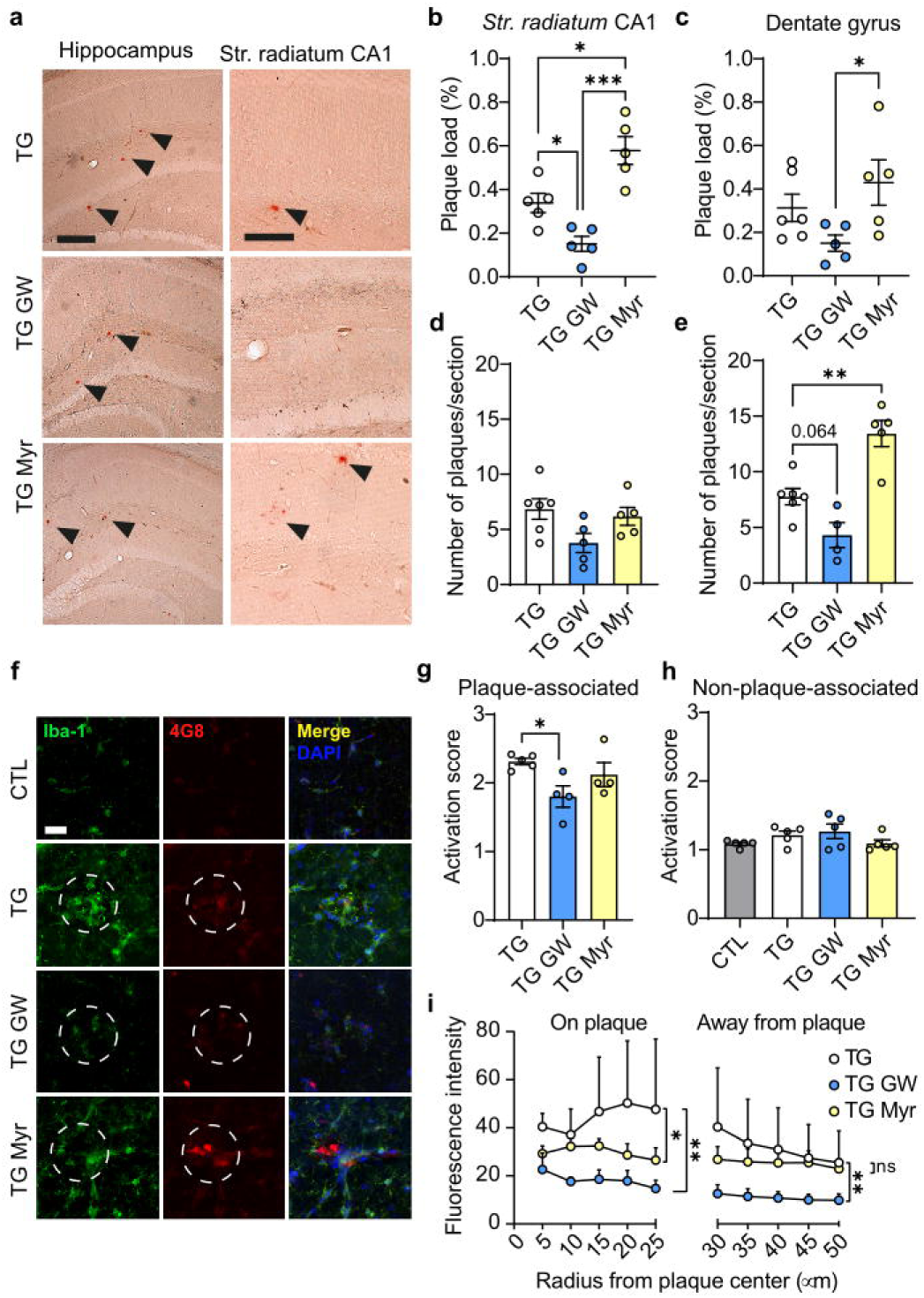
Amyloid plaques and microglial morphology. **a.** Microphotographs of Congo red-stained hippocampal sections. The left column shows images of the hippocampus including the dentate gyrus. Scale bar=200 µm. The right column shows images of the *stratum radiatum* under CA1. Scale bar=100 µm. Arrowheads indicate Congo red-stained amyloid plaques. **b and c.** Quantification of the mean plaque load per mice (% area covered by plaques) in the *stratum radiatum* under CA1 and in the dentate gyrus, respectively (n=5-6 mice per group). **d and e.** Number of plaques/section per mice in the *stratum radiatum* under CA1 and in the dentate gyrus (n=4-6 mice per group). **f.** Confocal images of Iba1 (microglia; green) and 4G8 (amyloid-β; red) stained hippocampal sections (*stratum radiatum* under CA1). Dotted circles indicate amyloid plaques and the peri-plaque area. Scale bar=20 µm. **g and h.** Morphological activation score for microglia associated or not associated to amyloid plaques, respectively (n=4-5 mice per group). **i.** Radial profile analysis of Iba1 staining at amyloid plaques (5-25 µm radiuses) and away from plaques (30 to 50 µm radiuses). Each dot represents the mean fluorescence intensity for each group and radius (n=4-5 mice per group). * P < 0.05, ** P < 0.01, *** P < 0.005, **** P < 0.001.

### nSMase Inhibition with GW4869 Attenuates Microglial Activation in PDAPP-J20 Transgenic Mice

To evaluate the impact of ceramide pathway inhibition on microglial responses in the context of amyloid pathology, we performed a double immunofluorescence for Iba1 and Aβ (4G8 antibody) on hippocampal sections (fig. 3f). To quantify the morphological activation of microglia, we classified Iba1-stained cells in the *stratum radiatum* under CA1 using an activation score. In the analysis of plaque-associated microglia (up to 25 µm from the plaque center), we found differences between groups (one-way ANOVA F (2,10) = 4.125 P < 0.05; fig. 2g) with the TG GW group showing a lower score than vehicle-treated TG (Dunnet’s P < 0.05) while The TG Myr group did not differ from TG. The activation score of non-plaque-associated microglia did not differ between groups (fig. 2h). To further characterize Iba1-stained microglia, we performed a radial profile analysis, setting the reference point on the center of each plaque and measuring the Iba1 fluorescence every 5 µm up to a radius of 50 µm. Analysis by region (5-25 µm for on-plaque microglia, 30-50 µm for microglia away from plaque) showed a treatment effect (RM ANOVA P < 0.001 for every region; fig. 2i). Treatment with GW induced a decrease at both regions (Dunnet’s P < 0.01 at each region vs. TG). Myriocin treatment was associated with decreased fluorescence intensity at the on-plaque region (P < 0.05 vs. TG) but did not differ from vehicle-treated TG at the farthest region. These results demonstrate that selective nSMase inhibition globally attenuates morphological microglial activation, whereas SPT inhibition with myriocin produces only a partial reduction in microglial reactivity restricted to the immediate vicinity of amyloid plaques.

**Figure 3.**
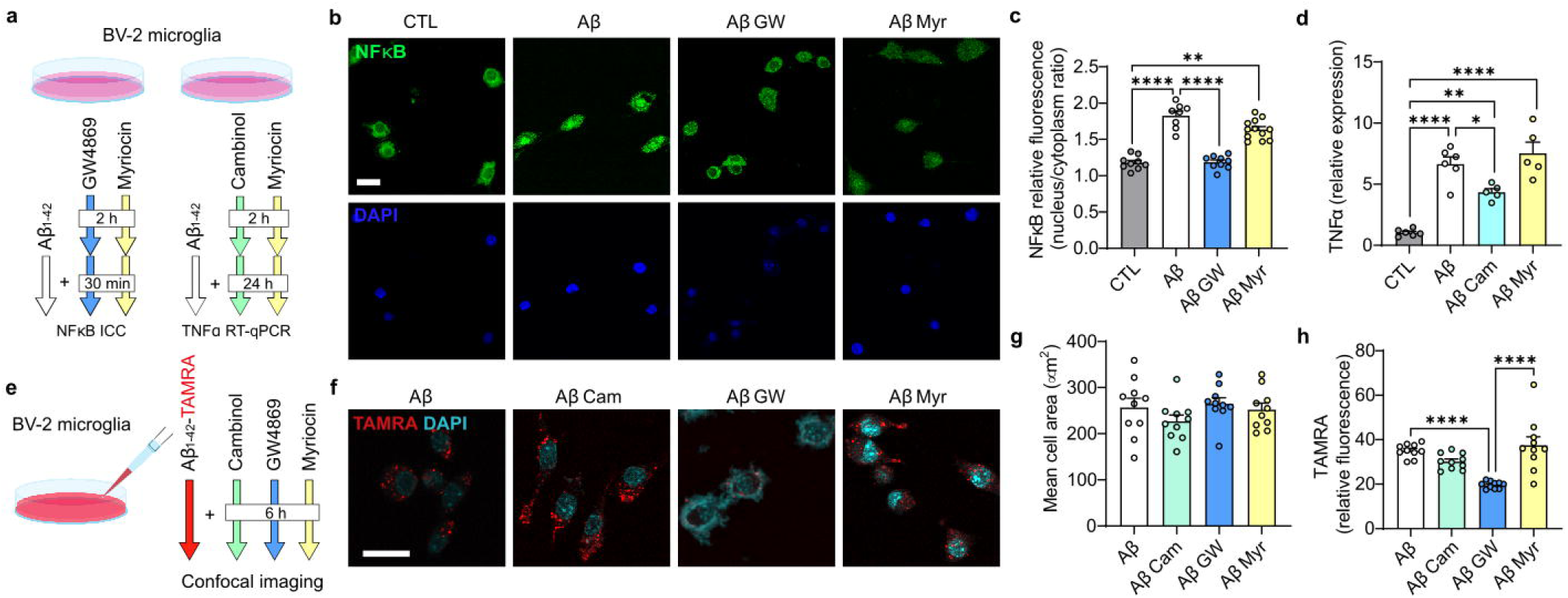
*In vitro* mechanisms in Aβ-challenged BV-2 microglia. **a.** Experimental scheme for *in vitro* treatments. Ceramide synthesis inhibitors GW4869 and myriocin (Myr) were applied to microglia for 2.5 h with fibrillized Aβ_1-42_ on the last 30 min to measure NFkB nuclear translocation. To obtain mRNA for TNF-α RT-qPCR microglia was exposed to fibrillized Aβ_1-42_ and inhibitors cambinol (Cam) and Myr for 24 h (a 2-h pretreatment with inhibitors was applied). **b.** NFκB p65 nuclear translocation was evaluated by immunofluorescence. Scale bar=20 µm. **c.** Quantification of NFκB translocation by the nuclear/cytoplasm fluorescence ratio (n=7-10 confocal fields with 5-10 cells nested per field; experiment repeated in duplicate). **d.** Expression of TNFα was evaluated by RT-qPCR (n=5-6 technical replicates per group from 3 independent experiments). **e.** Experimental scheme for evaluating amyloid internalization capacity. Microglia was incubated with TAMRA-Aβ1-42 and inhibitors Cam, GW and Myr for 6 h. **f.** Cells were imaged *via* confocal microscopy. Scale bar=20 µm. **g.** Mean cell area was measured to rule out an effect of treatments on cell size. **h.** Intracellular fluorescence intensity for TAMRA was measured to detect amyloid internalization (n for area and TAMRA intensity = 10 cells per group from 3 technical replicates). * P < 0.05, ** P < 0.01, **** P < 0.001.

### Selective nSMase Blockade Reduces Microglial Reactivity and Intracellular Aβ Accumulation in Fibrillized Aβ-Challenged BV-2 Microglia In Vitro

To elucidate the molecular mechanisms underlying the effects of ceramide synthesis pathway inhibition on microglial responses to amyloid pathology, we utilized the BV-2 murine microglial cell line exposed to fibrillized Aβ₁₋₄₂ (fAβ₁₋₄₂) for 1, 6 or 24 h as an *in vitro* model. Cells were treated with GW4869, Myr, or cambinol (Cam; alternative nSMase inhibitor; fig. 3a). Fibrillar Aβ₁₋₄₂ exposure induced nuclear translocation of NFκB (p65 subunit; quantified *via* immunofluorescence; fig. 3b), a key driver of pro-inflammatory gene expression, evaluated by the nucleus/cytoplasm fluorescence intensity ratio (nested one-way ANOVA F (3,31) = 19.31; Tukey’s P < 0.001 Aβ vs. CTL; fig. 3c). GW4869 treatment yielded levels similar to those of CTL cells and lower than the Aβ group (P < 0.001 Aβ GW vs. Aβ). On the other hand, Myr showed minimal or no effect. Consistent with NFκB modulation, qRT-PCR analysis revealed that fAβ₁₋₄₂ markedly upregulated TNFα mRNA (one-way ANOVA F (3,18) = 31.21 P < 0.0001; Tukey’s P < 0.0001 CTL vs. Aβ), a pro-inflammatory cytokine and nSMase activator. Treatment with the nSMase inhibitor Cam blunted this induction compared to vehicle-treated fAβ-exposed cells (P < 0.05; fig. 3d), though the expression did not reach control levels (P < 0.001 Aβ Cam vs CTL). TNFα expression in the Aβ Myr cells did not differ from vehicle-treated Aβ cells. Phagocytic capacity was assessed incubating BV-2 microglia with fluorescently labeled fAβ₁₋₄₂ (TAMRA-Aβ₁₋₄₂; 0.05 μM) for 6 h (fig. 3e). Intracellular incorporation was quantified by fluorescence intensity (*via* confocal fluorescence microscopy; fig. 3e). One-way ANOVA showed no significant differences in cell area across treatments, confirming comparable cell viability and size (F (3,36) = 1.153 P = 0.341; fig. 3g). However, a marked effect was observed for intracellular TAMRA-Aβ fluorescence intensity (one-way ANOVA F (3,36) = 14.95 P < 0.001, fig. 3h). GW4869 treatment reduced the intracellular signal relative to vehicle-treated fAβ-exposed cells (Tukey’s P < 0.001). Notably, this reduction was not replicated by Myr or Cam, which yielded levels similar (TG Cam) or higher (TG Myr) to those of vehicle-treated cells.

## Discussion

Collectively, our findings demonstrate that pharmacological modulation of ceramide metabolism exerts pathway-specific effects on key pathological hallmarks of AD. These differential responses highlight the distinct contributions of sphingomyelin hydrolysis *versus de novo* pathways to ceramide-driven neuroinflammation, amyloid progression, and cognitive decline, underscoring the therapeutic potential of targeted nSMase inhibition in AD.

In the PDAPP-J20 model, inhibition of *de novo* ceramide synthesis *via* SPT with myriocin (Myr) exacerbated anxiety-like thigmotaxis in the open field and impaired spatial learning recovery in the Barnes maze, while modestly increasing hippocampal amyloid plaque load. These adverse effects align with reports of Myr’s paradoxical outcomes in the mammalian CNS, including elevated brain ceramides in non-diabetic control ruminants (Davis et al., 2021) and failure to mitigate pathology in alcohol-related neuroinflammation in rats (Homans et al., 2022), likely due to compensatory sphingolipid rerouting or disruption of protective lipid homeostasis (Pan et al., 2023). Importantly, the detrimental effects of Myr may stem not only from central actions but also from the systemic inhibition of SPT following i.p. administration, which disrupts peripheral ceramide homeostasis and may indirectly exacerbate neuroinflammation via peripheral-to-central signaling

In contrast, nSMase inhibition with GW4869 produced clear therapeutic benefits: restored spatial learning, reduced hippocampal neurodegeneration (increased neuronal density and layer thickness), decreased amyloid plaque burden in the *stratum radiatum*, and inducing a widespread decrease in microglial activation, as shown by lower Iba1 optical density both in peri-plaque areas and in regions distant from plaques. These findings are consistent with prior studies showing that GW4869 reduces Aβ pathology and cognitive deficits in APP NL-F and 5XFAD models (particularly in females) (Mowry et al., 2023), lowers ceramide-enriched exosome release, preserves synaptic proteins, and modulates AMPA/NMDA receptor composition in hippocampus (Dinkins et al., 2014). While consistent with prior studies, our results provide novel evidence of neuroprotection at the structural level and a microglial phenotype dependent on its localization.

Mechanistically, nSMase inhibition likely reduces amyloid pathology through dual actions: decreased production (*via* disrupted ceramide-dependent exosome biogenesis and propagation of Aβ seeds) and enhanced degradation (Dinkins et al., 2014; Mowry et al., 2023; Tohumeken et al., 2023). *In vivo*, plaque-associated microglial recruitment suggests improved targeted phagocytosis or containment. In Aβ-challenged microglia, nSMase inhibition suppressed NFκB p65 nuclear translocation and downstream TNF-α transcription, interrupting the ceramide-TNFα-nSMase positive feedback loop. The marked reduction in intracellular TAMRA-Aβ fluorescence, without changes in cell area or uptake, points to enhanced lysosomal or endolysosomal degradation rather than a mere reduction in phagocytic uptake. These cellular effects align with the known role of nSMase-derived ceramides in modulating endosomal trafficking (Pavlic et al., 2023) and exosome biogenesis (Zhu et al., 2021), processes that propagate both inflammation and Aβ seeds.

In our *in vitro* experiments, we used both GW4869 and cambinol to inhibit nSMase. These chemically distinct inhibitors act through different mechanisms (Dinkins et al., 2014; Figuera-Losada et al., 2015; Mowry et al., 2023; Vinuesa et al., 2019). Although cambinol has limited pharmacological stability and is unsuitable for *in vivo* use (Kumar et al., 2019), both compounds yielded consistent beneficial effects. Thus, nSMase inhibition exerts protective actions against Aβ-induced microglial dysfunction regardless of the mechanism.

A limitation of the *in vitro* work is reliance on BV-2 cells, which exhibit blunted TGF-β signaling, reduced chemotaxis, and lower reactivity compared to primary microglia, potentially underestimating full glial responses *in vivo* (He et al., 2018; Luan et al., 2022). Nonetheless, the anti-inflammatory profile of GW4869 in microglia aligns with reports of reduced cytokine release and tau propagation following nSMase inhibition (Mowry et al., 2023). A limitation of our study is that only female PDAPP-J20 mice were used. This choice was made because female mice in this model develop more hippocampal neurodegeneration and glial reactivity at 8 months of age compared to age-matched males, as previously reported using PDAPP-J20 and other APP transgenic models (Clinton et al., 2007; Dhungana et al., 2023; Gregosa et al., 2019). This approach reduces variability and increases statistical power while aligning with ethical principles of minimizing animal use. However, it restricts the generalizability of our findings. Increasing evidence demonstrates significant sex differences in Alzheimer’s disease, both in humans and mouse models, including dimorphic microglial responses, amyloid clearance rates, and therapeutic responses to anti-inflammatory and lipid-modulating interventions (Reed and Keller-Norrell, 2023). Therefore, future studies should include both sexes to determine whether nSMase inhibition exerts similar neuroprotective and anti-amyloid effects in male PDAPP-J20 mice.

Neutral SMase emerges as a promising AD target due to its role in ceramide-enriched exosome biogenesis, which facilitates dissemination of inflammatory signals and pathological proteins (Tallon et al., 2021). Preliminary evidence from our lab supports ceramide modulation of small EVs release in amyloid-challenged glial cells, warranting further investigation into EV-targeted therapies. Direct neuronal effects of ceramides (e.g., apoptosis induction, Aβ stabilization) further justify pathway-specific inhibition over broad ceramide reduction (Baloni et al., 2022; Jazvinšćak Jembrek et al., 2015). Alternative approaches, such as CERT modulation or other nSMase inhibitors, merit exploration to optimize therapeutic selectivity (Crivelli et al., 2021).

In conclusion, by targeting nSMase, we simultaneously addressed multiple AD hallmarks (neuronal loss, amyloid burden, and chronic neuroinflammation) highlighting its potential as a disease-modifying strategy. Unlike broad ceramide-lowering approaches, selective nSMase inhibition preserves *de novo* sphingolipid synthesis necessary for cellular homeostasis. These results highlight the need for pathway-specific lipid-targeted strategies in AD and suggest future studies should examine sex differences, long-term EV dynamics, combination therapies, and translation to human iPSC-derived models or clinical cohorts.

