## Supplementary material for "Differential Effects of Neutral Sphingomyelinase and Serine Palmitoyltransferase Inhibitors on Alzheimer’s-Like Neuropathology"

**Supplemental material for Bellotto et al.**

| **Gene** | **Forward Primer (5′ → 3′)** | **Reverse Primer (5′ → 3′)** | **Cycling Protocol** |
| --- | --- | --- | --- |
| GAPDH | GACGGCCGCATCTTCTTGT | ACCGACCTTCACCATTTTGTCT | 95º10’  (95º15”; 60º60”) × 40 |
| TNFα | GAAAAGCAAGCAGCCAACCA | CGGATCATGCTTTCTGTGCTC | 95º10’  (95º20"; 60º30”; 72º60”) × 40 |

**Supplementary table 1.** Primers used for TNFα and GAPDH in RT-qPCR and corresponding cycling protocols.

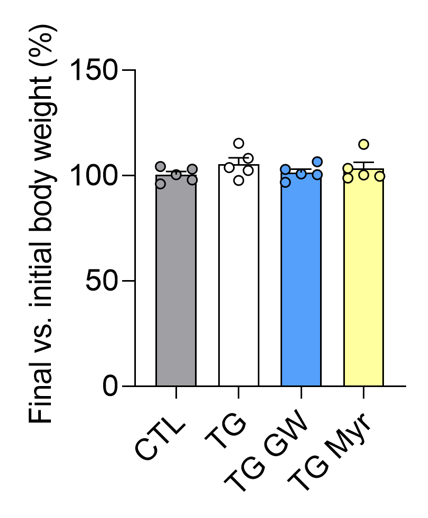

**Supplementary figure 1.** Comparison of final vs. initial body weight for each experimental group expressed as percentage. One-way ANOVA P = 0.47.

**a	b	c**

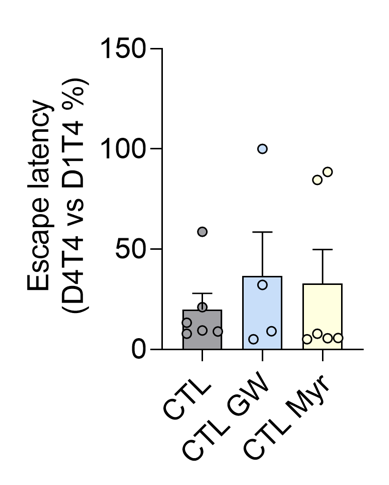

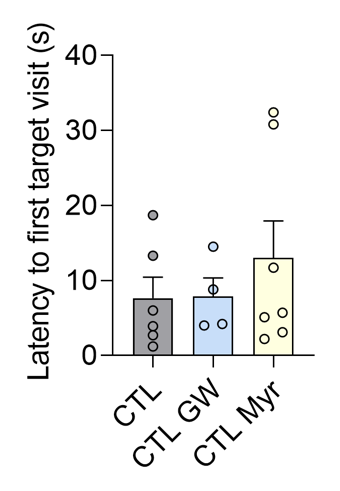

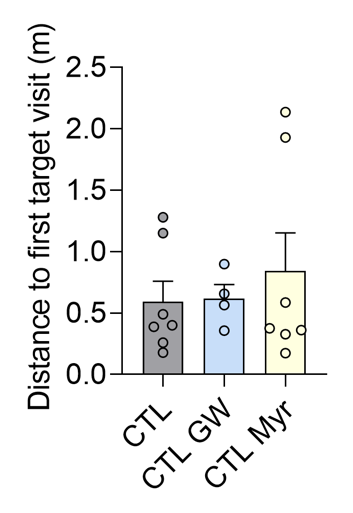

**Supplementary figure 2. a-c.** Recorded behavior in the Barnes maze for non-transgenic (CTL) mice with vehicle, GW or Myr treatments. **a.** Comparison of escape latencies between the 4^th^ trial of the 4^th^ day (D4T4) and the 4^th^ trial of the 1^st^ training day. Kruskal-Wallis P = 0.73. **b.** Latency to the first target visit on the 5^th^ day (test day). One-way ANOVA P = 0.47. **c.** Distance traveled to the first target visit on the 5^th^ day. Kruskal-Wallis P = 0.87.

**a	b	c**

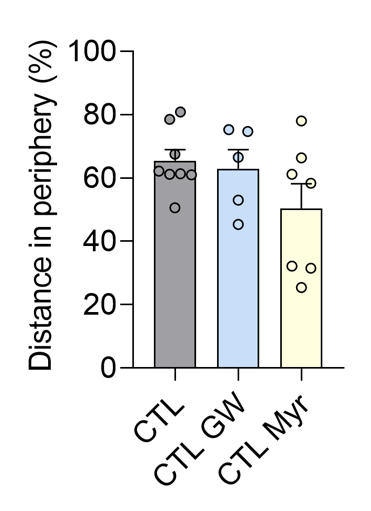

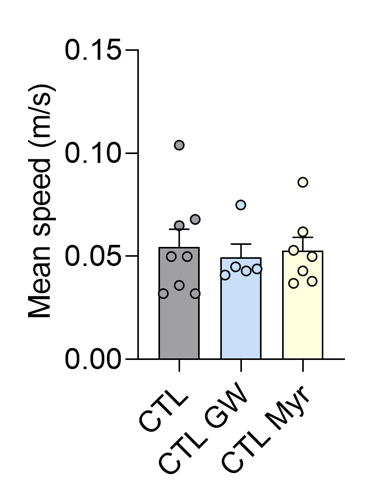

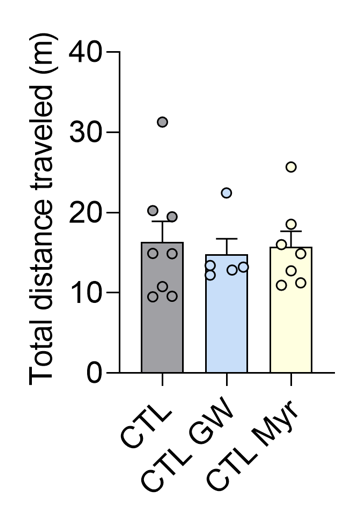

**Supplementary figure 3. a-c.** Recorded behavior in the open-field test for non-transgenic mice with vehicle, GW or Myr treatments. **a.** Percentage of distance traveled in the periphery. One-way ANOVA P = 0.16. **b.** Mean speed during the test. One-way ANOVA P = 0.91. **c.** Total distance traveled. Kruskal-Wallis P = 0.98.
